# A Platform for Semi-automatic Segmentation, Characteristic Shape Analysis and Simulation of Neuronal Nuclei

**DOI:** 10.64898/2026.11.24.768886

**Authors:** Shidan Javaheri, Athmane Benarous, Siger Ma, Thibaut Baguette, Aleksandra Bortel, Amir Shmuel

## Abstract

Segmenting neuronal somas in 3D brain images obtained through techniques such as tissue clearing and Light Sheet Fluorescence Microscopy promises to advance our understanding of healthy brains and the pathophysiological roots of neurological diseases. However, manual segmentation methods are often costly and time-consuming. Automating the segmentation through deep neural networks shows promise, but requires labeled ground-truth data that is also labor-intensive and expensive to obtain. To address these limitations, this work provides a platform within the open-source 3D image analysis software Vaa3D to efficiently a nd p recisely l abel a nd automatically segment neuronal somas. A binary image of segmented somas is generated using a region-growing algorithm applied on 3D somas marked and bounded by a sphere around their center of mass, guided by a local Otsu threshold that can be manually overridden. The platform further supports principal component analysis of the segmented somas, allowing them to be rotated about their center of mass and aligned, in order for their characteristic shape to be captured in a probabilistic model. Finally, the platform supports the simulation of soma segmentation: it probabilistically reorients, repositions and deforms segmented somas obtained from real data, generating synthetic ground truth data that can be used to train deep neural networks. The platform’s source code is released along with this paper, allowing future work to build on its initial architecture: https://github.com/orgs/ImagiNeuron/repositories.

## I. Introduction

THE advent of tissue clearing [1] and advanced 3D imaging techniques allows for the high-resolution imaging of the entire microscopic structure of the brain of animal models and part of the brain of humans, including isolating its neuronal somas. Analyzing and segmenting the terabytes of 3D image data of these somas allows the investigation of the structure and properties of neurons of the brain, advancing our understanding of the function of a healthy brain and the pathophysiological roots of neurological diseases. At this scale, the high volume of data is practically impossible to label manually, necessitating the automation of the segmentation process. Various efforts have shown the promise of deep neural networks and convolutional neural networks (CNN) in this regard [2]–[7]. However, the most accurate of these methods require ground truth data to train the machine learning models. Obtaining the ground truth data of segmented neurons is labor-intensive and time-consuming to obtain [2], [3], and software that supports it is often expensive and inaccessible to the broader research community.

LSFM is a 3D microscopy technique that uses a plane of light to image transparent tissue [8]. Used in conjunction with tissue clearing, it is an effective way to isolate and image neuronal somas in a whole brain of small animal models, or parts of a human brain. Another common imaging technique for analysis at this scale is Fluorescence Micro-Optical Sectioning Tomography (fMOST). This physical imaging technique is more common than LSFM in literature as the basis of the analysis of neuronal structures in the brain and in deep learning pipelines for the automated segmentation of neuronal somas [2], [4]–[6]. However, labeled ground truth data consisting of a binary segmentation of neuronal somas in the brain is rarely released for either of these imaging techniques. While the Big Neuron project [9] has focused on the automated reconstruction of singular neurons in Optical Microscopy Images and has released some ground truth data, none of it consists of binary labels for every voxel in a 3D image. Very recent work has focused on using a collaborative platform to quickly label the location of neuronal somas in the brain [10], successfully doing so for 180,000 somas in 122 different fMOST brains [11]. However, the work does not label somas in any brains imaged with LSFM, despite making reference to 4 LSFM datasets. While the project releases some of the data on 3D soma locations in the brain, this project and many other efforts in this area fail to provide access to binary segmentation of neuronal somas that can be used as ground truth data to train deep neural networks. There is a clear gap in accessible ways to efficiently and accurately obtain ground truth data of this nature, particularly for neuronal somas imaged with LSFM. To address this challenge, this work introduces a platform that aims to provide an accessible tool for fast and precise segmentation of neuronal somas in 3D microscopy images. In particular, its focus is on automatically segmenting neuronal somas that can be encapsulated by a sphere around their center of mass, without that sphere overlapping with other somas. In addition to supporting the correction of the automatic segmentation, the tool aims to support advanced visualization of segmentation results, including an overlay on the original image and a clear outline of the segmentation for its qualitative evaluation.

The platform further supports a tool to analyze the characteristic shape of somas with principal component analysis (PCA) on segmentation results. For comparison, intensity-weighted PCA on the original soma intensities is also supported, and both results can be visualized. PCA results allow all segmented somas to be aligned at their center of mass and along their principal components, to obtain and visualize a probabilistic model of the 3D shape of a neuronal soma. Finally, the platform supports data simulation, generating realistic 3D images of neuronal somas with synthetic shapes and intensities, as well as their corresponding binary segmentation. This simulation seeks to address the scarcity of available ground truth data, aiming to increase its quantity and diversity to train deep learning networks.

## II. Related Work

### A. Image Processing for Cell Segmentation

Given a 3D microscopy image of cells such as neuronal somas, factors such as intensity variations and noise make it difficult to segment and separate those somas from their background. To help accomplish this complex task without the need for manual labeling of individual voxels, image processing techniques have been developed to automatically or semi-automatically segment cells.

Some image processing techniques have been proven effective without manual labeling. One of these techniques is the watershed algorithm, which treats image intensity as a topographical map [12]. While relevant, the algorithm’s segmentation is prone to overflow from its intended soma regions, which is especially true in noisy images [13]. Due to the algorithm’s global approach and variation in image intensity in different parts of the imaged volume, the watershed algorithm often performs poorly in parts of the imaged volume. Alternatively, algorithms such as region-growing can use manually labeled seed points to initiate the segmentation of a cell. At its core, a region-growing algorithm iterates by recursively segmenting neighbors of the currently segmented voxels on the condition that they are below a certain intensity threshold. The most important element of region-growing is selecting an appropriate threshold to stop the expansion of the segmented volume and to accurately determine the 3D boundary of the cell body, separating it from the image background. There are various methods for selecting the optimal threshold. Choosing a global threshold for the whole image does not lead to an effective segmentation, as the intensity and boundary of every cell are different. Most cases require selecting a specific threshold for each cell body.

One approach to thresholding is called ‘iterative thresholding’ [14]. This method involves starting with a very high threshold for each cell body, and then successively lowering it, applying region growing with each threshold in sequence until the segmentation’s center of mass strays past a limit defined on its distance from its previous position. This limit is used as a heuristic for over-flooding. This approach is inefficient and does not always succeed, necessitating further heuristics to keep or discard a segmentation before over-flooding is detected [14]. However, its overall architecture and code is useful, and form a basis upon which part of our work is built, as discussed in Section III.

Research indicates the effectiveness of local Otsu thresholding to separate the voxels of a cell from the background [15]. This method uses a histogram to separate foreground from background voxels and seeks to choose an optimal threshold by minimizing intra-class variance and maximizing inter-class variance [15]. This approach was selected as the basis of the thresholding in our design.

### B. 3D Visualization-Assisted Analysis (Vaa3D)

3D Visualization-Assisted Analysis (Vaa3D) is an open-source software platform for the visualization and quantitative analysis of large-scale biological image datasets [16]–[18]. The platform supports viewing and labeling images in 2D and 3D. Labeling is supported by placing markers on cells in the image to contain each cell within a sphere around its center of mass. The platform also supports TeraFly, which provides Google Earth-like functionality that supports efficient navigation through terabytes of data. The data is loaded on the fly, and the resolution is adjusted as the user navigates through it at different levels of resolution [19]. TeraFly also comes with a tool called TeraConverter [20], that allows the automated conversion of 2D microscopy images taken at various slices in the z-axis into the 3D format supported by TeraFly. In addition to Terafly, Vaa3D also allows developers to add custom plugins, supporting the development of any additional functionality. It provides a strong basis to build robust and accessible pipelines to facilitate medical image analysis, particularly given its recent update [21].

### C. Deep Learning based Soma Segmentation

Deep learning and CNNs have been solutions on the rise for computer vision tasks such as biomedical image classification and segmentation [22], and they often outperform more traditional computational and machine-learning methods. However, they rely on the availability of large labeled datasets to prevent overfitting and improve their generalization. There is often limited biomedical image training data, and producing more well-annotated data labelled manually by human raters can be expensive and time-consuming [2], [4].

Several efforts have explored applying deep neural networks for segmenting somas from 3D microscopy data, by avoiding using a complete binary segmentation of the brain as ground truth data. The following brief overview of a few of these methods indicates that while they are somewhat successful, they are less effective than those trained with more extensive ground truth data. Most of the studies use fMOST 3D images of the brain, with no studies done on LSFM data to our knowledge.

Recent work [5] explores using deep neural networks to locate the centers of neuronal soma in fMOST images of the brain rather than explicitly segmenting them, given that manual labels of the centers of mass of somas are faster to obtain. The method uses a 2-stage neural network and a custom loss function to define the objective of correctly locating soma centers. While it achieves a high precision of 98%, its design ensures a low false positive rate, and it does not classify neuronal somas in dense brain areas, reflected i n i ts l ow F1 score of 64%.

Another study [6] explores what it refers to as a “weakly supervised” approach. Ground truth data consists of labels for each neuronal soma and a radius defining a s phere that completely envelopes each soma, labels which are also relatively easy to obtain. For training, a simple approach is used with no further refinement - v oxels a re c lassified as “foreground” within the sphere and “background” if they are far enough from all other spheres, with other voxels ignored. This approach leads to a precision of 78%, and a recall of 83%.

Another method [4] attempted to avoid using ground truth data altogether. Working with fMOST images of the brain, it uses a “Hessian analysis-based enhancement filter” t o automatically create weakly supervised labels for all voxels. A deep neural network is trained on this data to segment neuronal somas, with reported results with a high F1 score of 97%. However, our further investigation into the project’s code base revealed that the results may be misleading and not reproducible, as we found little evidence supporting the paper’s claims.

Methods that use a binary segmentation of somas as ground truth data have achieved better results. The UNET CNN architecture is well known for its high performance on medical image analysis [7]. Recent work [2] cites the promise of building on this architecture for soma segmentation applications, and adapts UNET to include multi-scale feature extraction to enhance finer d etails, a daptive w eighting t o m easure feature importance, and dynamic convolution to address the sparsity in neuronal soma data. It achieves an F1 score of 91.8%, an improvement compared to other UNET architectures it is compared to, including 3D UNET [23], SegNet [24], HRNET [25], Attention UNET [26], Res UNET [27], and RAUNET [28]. Despite its high performance, its code and ground truth data have not been made available.

A brief survey of methods for soma segmentation indicates the importance of obtaining a binary segmentation of neuronal somas in the brain as ground truth data. However, data of this nature is costly to obtain and rarely made publicly available, particularly for LSFM data. Our platform seeks to address these limitations by providing a means to efficiently o btain a binary segmentation of neuronal somas to be used as ground truth data, as well as to enhance its quantity and diversity through data augmentation and simulation.

### D. Data Augmentation

Data augmentation is essential for addressing the high cost of obtaining a large amount of labeled data necessary to build high-performing deep learning and CNN pipelines [29]. Data augmentation methods alter the original image with a well-defined operation and add it to the dataset on which the model is built, improving the data’s quality and diversity. This helps the model experience different scenarios and learn from them. Other image data augmentation techniques exist, including basic, deformable, and deep learning augmentation techniques [29]. Table I summarizes these data augmentation techniques and their categories.

**Table I.** Summary of data augmentation techniques and their categories [29]–[31].

| **Category** | **Technique** | **Description** |
|---|---|---|
| Basic | Geometric Transformations | Scaling, translating, rotating, reflecting, shearing, skewing |
|  | Cropping | Select a random patch from the image |
|  | Occlusion | Remove random patches from the image |
|  | Intensity Operations | Pixel/voxel manipulation; gamma correction; contrast enhancement |
|  | Noise Injection | Add Gaussian, uniform, or salt-and-pepper noise |
|  | Filtering | Apply kernels to sharpen, blur, or smooth the image |
|  | Combination | Combine two or more original images |
| Deformable | Displacement Field | Random value shifts each pixel/voxel |
|  | Spline Interpolation | Use piecewise polynomials to estimate values between data points |
|  | Image Registration | Align moving image to match a fixed image |
|  | Shape Models | Generate shape variability models across the dataset |
| Deep Learning-Based | GANs | Generate realistic images via adversarial learning |
|  | Feature Augmentation | Augment representations in feature space |

Deep learning approaches for segmentation tasks require image-label pairs. Data augmentation can generate a substantial amount of synthetic image-label pairs semi-automatically, using a limited number of well-labeled examples encompassing a diverse range of neuronal somas from different brain regions. However, it requires an initial dataset of labeled somas that can be used for data augmentation.

GAN-based methods have been used for data augmentation techniques to generate realistic synthetic image-label pairs suitable for image segmentation [29]. GAN-based data augmentation methods require a significant amount of initial labeled data for training [30], which poses practical challenges for datasets that contains raw image data but lack sufficient labeled examples.

Deformable data augmentation techniques can generate realistic synthetic image-label pairs through deformations with plausible parameters. They are especially effective for datasets with limited labeled training data to improve segmentation results through generating additional labeled, realistic data. Previous studies have used this data augmentation technique to improve their biomedical image segmentation performance [32]–[35].

## III. Method

### A. Dataset

We used one dataset consisting of LSFM images of a rat brain imaged at a magnification of 9.0X. The dimension of each voxel were 0.71*µ*m × 0.71*µ*m × 1.0*µ*m (x, y, z). All nuclei were segmented and simulated at their original intensities, without changing image intensity or contrast.

### B. Overview of Overall Architecture

The platform we built is designed and provided as a plugin to Vaa3D, making the most of this open-source platform’s visualization, navigation, labeling, and image processing features. We further ensure that the functions in our plugin are compatible with TeraFly, allowing for simple conversion of 2D slices of microscopy data into a 3D format, and the seamless navigation through and analysis of terabytes of data. We name our plugin Machine Intelligence for Neuron Detection and Segmentation (MINDS); all functions are accessible under the MINDS plugin.

Our platform uses Vaa3D’s built-in markers to provide semi-automatic segmentation of somas with minimal manual labeling, allowing users to isolate and contain a soma within a bounding sphere around its center of mass. These labels are faster to obtain than manually labeling each voxel. The platform provides the basis for a region-growing-based segmentation algorithm to automatically generate a binary segmented image, which is then also visualized in detail for qualitative analysis of results. The platform is thus built on the premise that it will only be able to label and process somas that can be isolated in this way, leaving extensions to overlapping somas in the brain for future work.

Deformation and shape modeling data augmentation techniques align well with our dataset, segmentation, and augmentation objectives. From a limited labeled dataset, statistical shape models can combine the available data to describe neuronal somas’ shape, position, and orientation variability with probabilistic models. To be able to simulate images of somas, we need to first build these probabilistic models, by analyzing the shape of somas.

The platform thus supports analysis of the characteristic shape of somas. When a binary segmented image of somas is generated, the outputs include files containing the PCA results on the 3D coordinates that each segmented soma occupies, along with a probabilistic model of a soma shape built from superimposing all the aligned segmented somas. These outputs can be visualized through separate functions provided by our plugin. Finally, the platform uses the segmentation files and analysis of soma shapes to generate realistic simulations of soma data and their corresponding binary segmentation. This feature provides data augmentation critical for building deep learning pipelines to completely automate the soma segmentation task. This plugin’s main workflow and use are depicted and described in Figure 1. The details of each function are outlined further during our presentation of results in Section IV-A.

**Fig. 1.**
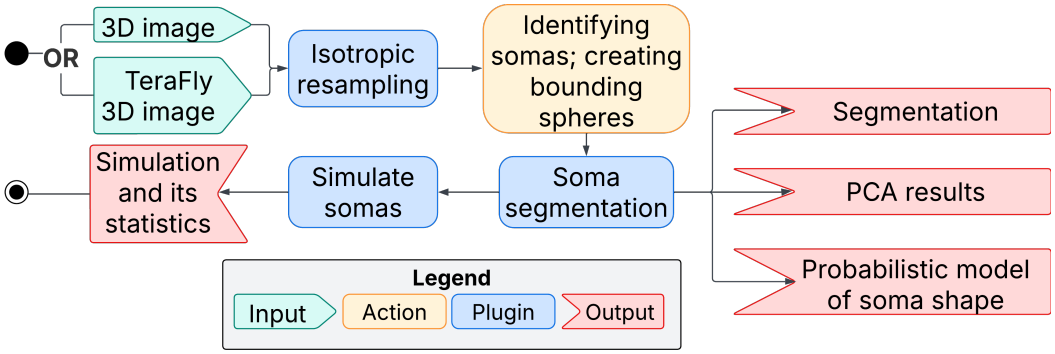
The main workflow and architecture of the segmentation and simulation pipeline. Once a user views an image in Vaa3D, either through TeraFly or by loading a 3D image into Vaa3D directly, the plugin’s functions are accessible. The isotropic correction function is used to obtain isotropic dimensions of a voxel for 3D visualization and calculations. The original dimensions are often not isotropic; they are determined by the resolution of the microscopy technique and the separation between subsequent Z slices. The user can identify somas and bound them within spheres of varying radii centered at the soma’s center of mass. Once a sufficient number of somas have been labeled, the image can be segmented with the soma segmentation function. This function produces an output of the binary segmented image of each manually identified soma, the results of PCA on its segmented shape, and the probabilistic model of soma shape. With these results, simulation of soma data can be generated with a custom number of synthetic somas, using various parameters that control the nature of the simulation.

**Fig. 2.**
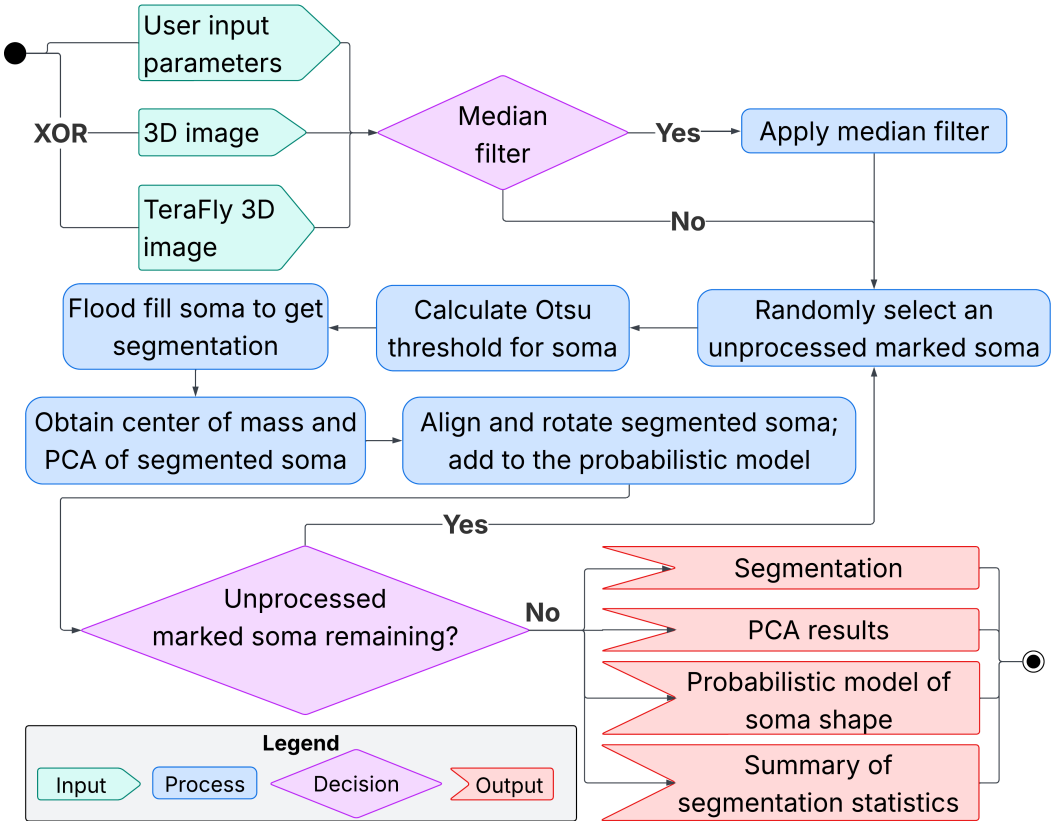
The main workflow and architecture for soma segmentation, using local Otsu thresholding applied to each soma

Our platform, therefore, has four key features: the semi-automated segmentation of somas, PCA of segmented somas, probabilistic modeling of soma shape, and soma simulation. The method used to achieve each of these functionalities is discussed in detail in the following sections. During processing and analysis, the outputs of each function are always automatically saved. A different approach is used to save files processed with and without TeraFly, and both are clearly documented.

### C. Segmenting Somas

Our solution for semi-automated soma segmentation uses an existing Vaa3D plugin as a basis, drawing from its implementation of region growing to segment cells [14], [36]. We vastly improve on the iterative approach used in [14], [36] by using local Otsu thresholding to find the optimal threshold to segment each soma - as highlighted and discussed in Section II. This function of our plugin is designed to run once the user has isolated all the somas they want to segment within a sphere around their center of mass, using the markers in Vaa3D.

The soma segmentation function allows the user to preprocess the 3D image using an efficient histogram-based 3D median filter, using a user-defined window size. This filtering approach aims to reduce salt-and-pepper noise without blurring the edges of somas [37], thus, it is ideal for our segmentation task. Once the segmentation process is initiated, a 3D region-growing algorithm starts from the center of each soma, which have been labeled by the Vaa3D markers. Along with its center, each marker has a radius defining the volume of the bounding sphere, which, by default, also defines the maximal segmentation volume. The user has the option to select one of three methods to determine the optimal intensity threshold used to flood soma voxels. The first two options are provided for comparison, but the optimal one is the third method:

1. Iterative Thresholding: This method was initially provided by the reference plugin [14], and it determines the threshold for each cell iteratively. This iterative thresholding and its associated hyperparameters were left as part of the plugin for reference and comparison, but their use is not recommended.
2. Global Otsu Thresholding: This method determines an optimal threshold to separate the foreground and background of the whole image after being given a histogram of the entire 3D image. This method is also for reference and shows the weakness of using a global thresholding method, which is not recommended.
3. Local Otsu Thresholding: Instead of considering the whole 3D image, this method determines an Otsu threshold specific to the spherical volume defined by the Vaa3D marker around each soma. This allows each calculated threshold to better reflect each soma’s local intensities. This specificity feature of the local Otsu threshold produces the best results and is the method provided by default.

By default, the behavior of the segmentation algorithm is to constrain the segmented region of each soma to remain within the spherical volume defined by each soma marker, consistent with Vaa3D’s built-in markers function, which we used to identify somas. However, the user can disable the “marker constraint” option in the segmentation functions configuration. Disabling this option was added for flexibility, but is not recommended.

After completing the function, the segmentation results can be adjusted and corrected to increase accuracy. The intensity threshold value determined for each segmented soma using the user-selected thresholding method is automatically saved into the comments field of the Vaa3D marker. For example, a soma with an optimal threshold of 30 would have the following comment: “threshold:30”. Should the user notice an under-flooded or over-flooded soma segmentation through visual inspection, they can manually change the numerical threshold value stored in the marker’s comment. Selecting the “Manual Thresholding” option and rerunning the segmentation plugin will enforce the new threshold constraint for the specific soma. This allows the user to direct correction efforts of specific somas without affecting others.

When the segmentation processing step within our plugin is complete, the segmentation visualization is automatically opened in a new image window. This image contains three channels, each presented with a different color. The first channel contains the original image data, the second contains the segmented binary image data, and the third contains the gradient of the binary intensity of the segmented image. All three channels can have their color customized, such that, for instance, the original image is shown in grayscale, the segmentation is in red, and the outline is in cyan. The channels can also be individually hidden if desired. This new image can also be linked to the original image to show the same location in the 3D image as the user scrolls through one of the axes. Finally, the image can also be opened in the 3D viewer to have a global view of the segmentation. The image gradient is computed using a Sobel filter [38] applied in three dimensions, implemented with the OpenCV library for C++ [39]. An additional step of non-maximum suppression is applied to get a thinner edge, allowing the user to see the shape of the segmentation more easily on top of the original image data. The results of this function can be are visualized in Section IV.

### D. Principal Component Analysis of Soma’s shape

As one of its steps, the soma segmentation pipeline includes 3D PCA of the shapes of the volumes of segmented neuronal somas. PCA is performed on the coordinates of the neuronal soma’s voxels to capture their shape, after the center of mass of the soma has been calculated. The first principal component *PC*_1_ describes the orientation of the shape axis with the highest variance, representing the major axis of anisotropy. The second principal component *PC*_2_ is the second longest axis orthogonal to the first, and the third principal component *PC*_3_ is the third longest axis orthogonal to the two others. These three components summarize the spatial variance and anisotropy of the soma’s shape around its center of mass to its boundary [14]. They are captured in a shape feature vector composed of 3 normalized eigenvectors *eig*_*n*_ and their 3 eigenvalues 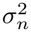, where *n* ∈{1, 2, 3} . The direction of each principal component is thus represented by its eigenvector, and its length is captured in its eigenvalue.

The statistics gathered for each neuronal soma through PCA on the binary segmented image focus solely on spatial and shape information. These parameters serve to create a probabilistic model of soma shapes and the simulation processes. In addition, the platform supports intensity-weighted PCA on the unsegmented image of each soma. This makes it possible to record information on voxel intensity to characterize the soma’s intensity, which can be used for simulation.

The PCA data can be visualized using our plugin’s “visualize PCA” function. Through this, a new color image window is opened with the original image and, for each segmented soma, a representation of its three principal components. The longest principal component is colored green, the second longest red, and the third cyan. In this case, the colors cannot be changed. The new image will be opened with the default 2D viewer of Vaa3D, but due to the nature of the visualization, in which each principal component is a thin 3D vector, it is preferred to view it in 3D. The eigenvectors are drawn as lines starting from the center of mass using Bresenham’s line algorithm [40], which allows fine l ine r epresentations i n a low-pixel-density scenario. To obtain an accurate estimate of the length of each principal component in 3D, the full width at half maximum measure is used [41], and the underlying distribution along each component is assumed to be gaussian. For the principal component represented by the eigenvector *eig*_*n*_ and the eigenvalue 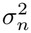, this value can be calculated as 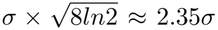. These results are further explored and visualized in Section IV.

### E. Probabilistic Model of Soma Shapes

The soma’s center of mass, as well as the eigenvectors *eig*_1_, *eig*_2_, and *eig*_3_ defining the principal components of the binary volume from the segmentation of each soma, allow us to document a soma’s position and its orientation in 3D space in the analyzed volume. With this information, we can place the center of mass at the origin of a 3D Cartesian coordinate system. We can then rotate the segmented soma about its center of mass to align its principal components with the Cartesian axes, such that *eig*_1_ aligns with the y-axis, *eig*_2_ aligns with the x-axis, and *eig*_3_ aligns with the z-axis. Once all the somas have been aligned to this coordinate system, we can sum all the binary soma volumes to get a count value for each voxel, indicating the number of somas each voxel is a part of. By dividing by the total count of somas, we obtain a probabilistic model that characterizes soma shape in the volume we analyzed.

A simple heuristic is used to check whether the segmentation failed. If, after rotating the binary segmentation of a soma about its center of mass, the value of the voxel at the center of mass is 0 (i.e, not part of a soma), the soma is flagged as having poor segmentation, and a notification is displayed to the user. The output in the Vaa3D terminal displays a slice through the center of the binary volume of the rotated soma, allowing the poor segmentation to be visualized. The user can then adjust the marker of the corresponding soma to address the issue before segmenting again.

The rotation of the soma from a source image into a target image where its principal components are aligned with the Cartesian axes is approached as follows. Once a single soma is segmented, its segmented volume can be extracted to create a source image, in which the soma’s principal components are not aligned with the Cartesian axes. This source image is binary, where the voxels that are part of the soma have a value of 1. The coordinates of every point in the volume of the source image relative to the soma’s center of mass can be calculated and used in all future operations. A rotation matrix *A* can then be defined, whose columns *a*_1_, *a*_2_, *a*_3_ are the eigenvectors representing each principal component of the soma, *a*_1_ = *eig*_2_, *a*_2_ = *eig*_1_, *a*_3_ = *eig*_3_, respectively. Multiplying by *A*^*T*^ maps coordinates in the source image to the target, and multiplying by *A* maps coordinates in the target image to the source.

This rotation operation is prone to numerical errors that need to be mitigated. Errors may arise because the principal components defining a soma’s orientation support floating point precision, and can be in any three orthogonal directions. However, the volume in which somas are represented is discrete, with only integer coordinates. This means that any mapping between the coordinates of the source and target matrices using matrix *A* maps integer coordinates to floating points, which then need to be rounded to be integers in order to visualize the target image, creating numerical imprecision. A simple trilinear interpolation method was employed to address numerical errors when generating the target image. The target image is generated by iterating over every voxel to ensure that every voxel has a value. Each coordinate in the target image is mapped to the source image by multiplication with the matrix *A*. Since the resulting coordinate in the source image is a floating point, we use trilinear interpolation along all three axes to obtain the value for the target voxel in the target image. The target image generated thus contains a 3D volume of floating points with floating values between 0 and 1 generated by trilinear interpolation, describing the soma’s shape, which is now aligned with the Cartesian axes.

Each aligned soma is added sequentially to construct a model that probabilistically describes the overall soma shape. Dividing the value of all voxels by the value at the center of mass (which is equal to the number of somas aligned and included in the probabalistic model) gives the probability of each voxel in the 3D volume being part of a soma. The probabilistic model is saved as a binary file in a simple format: the first 24 bytes contain three 64-bit integers representing the image’s x, y, and z dimensions. The rest of the file contains double-precision floating-point values representing the volume of the probabilistic model of soma shape.

We can visualize the probabilistic model’s results using the plugin’s “visualize probabilistic model” function, which loads a user-selected probabilistic model in a new image window and represents it as a heatmap. This will open by default in Vaa3D’s 2D viewer, but can also be opened in a 3D view. A legend will also open in a separate window to present the color mapping of the heatmap. These corresponding results are presented in Section IV.

### F. Simulation of Somas

The proposed platform also allows augmentation of existing ground truth data through statistical shape models to increase the quantity and diversity of realistic image-label pairs. Figure 3 illustrates the methodology of the simulation pipeline. The simulation takes as input the original image as input, its segmentation, the probability model of soma shape, and the characteristics of the neuronal somas, including each soma’s ID, marker coordinates and radius, center of mass, and principal components. We obtain the normal distribution of the soma’s position and orientation from each soma’s center of mass and principal components. Somas that have already been segmented are randomly selected to be used as the basis of a simulated soma, and then deformed, reoriented, their intensities randomly offset by a small constant, and then placed into two synthetic images: an image containing simulated somas with synthetic intensities, as well as its corresponding binary segmentation. Once all synthetic somas and their segmentation are generated, the two images are output. The simulated soma’s shape is created by probabilistically deforming the original soma’s shape using the probability model of soma shape, and its intensity is also simulated through probabilistic analysis.

**Fig. 3.**
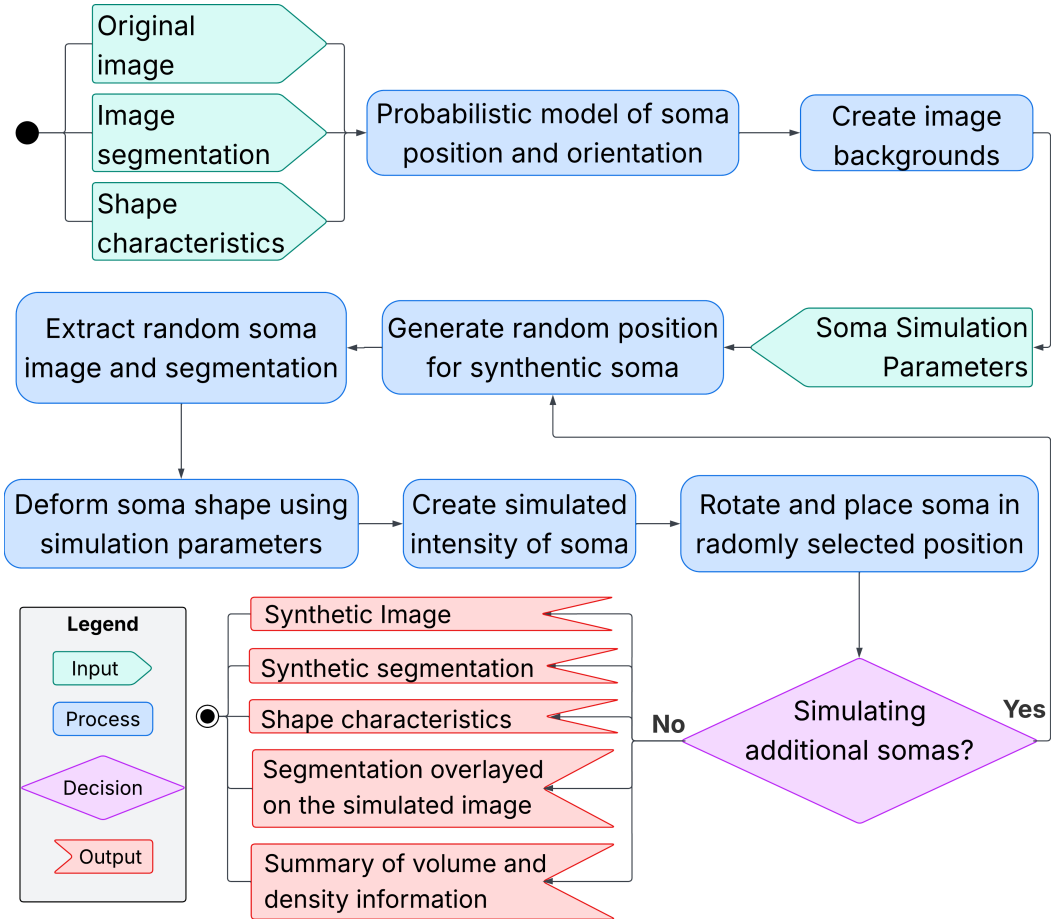
Workflow and architecture of the soma simulation pipeline

The simulation first g enerates t wo s ynthetic backgrounds, one for the synthetic image and one for its binary segmentation. To capture the complex background intensity variations observed in real data, the realistic background is not uniformly defined g lobally f or t he i mage. I nstead, l ocal i ntensity variations in the image are accounted for by dividing the image volume into 16 smaller, local 3D chunks. An intensity histogram is computed within each chunk, and an Otsu threshold [42] is determined and applied to each chunk to separate soma voxels from their background. Gaussian modeling, which has been used in a previous study to synthesize neuron images [43], is then calculated using the mean and standard deviation of the intensity values of the identified background voxels from each chunk. To ensure a coherent final b ackground i mage, instead of simply obtaining voxel intensities from the Gaussian distribution of each chunk, we perform gradual interpolation of the statistical parameters of surrounding chunks using distance-based weighting, which has been used to stitch microscopy imagery in a previous study [44]. This allows us to calculate Gaussian statistics for each voxel by considering its distance to each neighboring chunk up to a set blending radius, which we chose to set as a relative measure, at one chunk. That weight is also set to fall off quadratically as the distance decreases, to preserve a local chunk’s intensities. Once normal distributions based on the blended Gaussian parameters are calculated for each voxel, we generate its pixel value using the Box-Muller transform [45]. This gives us the background for the synthetic image of neuron somas.

The simulation pipeline generates synthetic images of neuronal somas, along with their binary segmentation, as follows. The user can input various values that define how the simulation will work, including the number of simulated somas to create in the generated image, and then starts the simulation. The method first l oads t he p robabilistic m odel of soma shape (soma shape model) from where it was saved during segmentation, and calculates other important normal probability distributions for this area of the brain. This includes the distributions of the eigenvectors that define t he principal components of soma shapes (eigen vector model), as well as the distribution of the positions of somas (position model). This probabilistic data can then be used to generate each synthetic soma iteratively.

To generate each synthetic soma, the method first randomly selects a soma from the original image that has already been segmented. The shape of this soma is then deformed along the direction of its eigen vectors, using a user-defined deformation factor that is scaled based on the distance of each voxel from the soma’s center. The deformed shape is then sampled against the probability model, and only voxels above a certain probability threshold are kept. The same operation is applied to the soma’s segmentation, so that they match. The soma is then rotated to a random orientation according to eigenvectors generated from the eigen vector model.

The three eigenvectors need to be orthogonal to each other for a single soma. The method employs QR decomposition with Householder Reflections to obtain three orthogonal vectors, ensuring numerical stability [46], done using the Eigen library in C++ [47]. The pipeline places these rotated synthetic soma images and their corresponding segmentation within their respective generated background. Finally, the intensity of the simulated soma is created by adding ± 10% of the range of intensities in the soma to every voxel, randomly making the soma slightly more or less bright.

The simulation pipeline finally saves the generated image and the segmentation image-label pair separately. The pipeline also saves its byproducts, including marker information and PCA data on synthetic neuronal somas. The pipeline displays the results in a window showing the segmentation overlayed on the image, with the same view as the simulation results. All files output by the simulation function are saved within a timestamped folder, allowing multiple simulations to be generated from the same segmented image. These results are discussed further in Section IV

There remains room for future work to improve limitations with the current simulation pipeline. The generated soma shapes and intensities were not thoroughly explored, but rather were generated with very simple probabilistic methods from existing somas segmented in the original image. Future work can improve the methods of deforming them based on the soma shape model, as well as better methods of creating synthetic intensities. Methods that could be explored include deformable data augmentation methods, such as spline interpolation [34].

## IV. Results

### A. Vaa3D Plugin

Our plugin for Vaa3D is called MINDS, and contains five main features, available as different entries within the plugin’s menu: isotropic correction, principal component analysis, soma segmentation, visualization, and soma simulation. The paragraphs below go through each functionality, briefly explaining what it does and how to use it. Details on how to use each functionality with images is documented in our code repository: https://github.com/orgs/ImagiNeuron/repositories.

#### 1) Isotropic Correction

The isotropic correction feature allows the user to input the dimensions of the image voxels along the three dimensions in micrometers to correct for anisotropic data. This affects any calculations that rely on accurate voxel dimensions, and 3D visualizations that use interpolations to show how the data should truly look.

#### 2) Principal Component Analysis

This function performs an intensity-weighted PCA of each labeled soma in the current volume. For each soma, the PCA data generated includes the soma’s ID, the x, y, z coordinates of the manual landmark, the radius of the sphere containing the soma, the x, y, z coordinates of the center of mass of the soma, the nine components of the three eigenvectors, and the three eigenvalues. All these results are automatically saved to a comma separated value (CSV) file that can be used for further analysis. The output of this function gives information about the intensity of labeled somas, and our plugin does not currently support a way of visualizing its results.

#### 3) Soma Segmentation

This feature uses the labels placed by the user to segment somas automatically. It produces a binary output image where each voxel shows whether the associated voxel in the input image was part of a neuronal soma. A pop-up window appears when selecting this option, asking the user to select the parameters for the analysis and whether or not to use pre-processing. By default, this function uses median filtering with a moving 3D window size of 3 voxels and local Otsu thresholding with flood filling, where the flooding is restricted to within the defined spherical marker. The processing starts when the user clicks the “run” button. Logging information will be printed to the terminal, and once the processing is finished, a pop-up window will open showing where the output files have been saved. A new image window will also be opened to show the results of the soma segmentation.

#### 4) Visualization

The plugin features two visualization functionalities: one for the PCA data (the output of the PCA function or the soma segmentation function) and one for the probabilistic model (the output of the soma segmentation function). When selecting the “visualize PCA” option, the user will be prompted to select the CSV file containing the data. Note that the original input image must first be opened. Then, the plugin will open the visualization. The “visualize probabilistic model” will prompt the user to select the probabilistic model file. Then, the plugin will open a new image window to present the probabilistic model as a heatmap.

#### 5) Soma Simulation

The soma simulation feature uses the method described in Section III-F to create new synthetic data. An image containing real data must first be opened to run this function, allowing the simulation to be based on it. Then, the user is required to input how many synthetic somas they wish to generate, and the data will be generated and automatically saved before being presented to the user using the 2D viewer, with the possibility to visualize it in 3D as well. Information about the resulting image volume, overall soma density, and soma density is calculated per image octant and presented to the user in a summary file .

### B. Segmentation

Due to a lack of available ground truth datasets for 3D brain microscopy, particularly for LSFM microscopy of neuronal somas, quantitatively evaluating our segmentation algorithm’s accuracy proved to be beyond the scope of our project. As a result, segmentation performance was primarily assessed through qualitative visual inspection using tools integrated into the Vaa3D plugin. Given the known drawbacks of global and iterative thresholding approaches, this qualitative analysis was done with the plugin’s default settings, using local Otsu thresholding.

Qualitative inspection using the aforementioned visualization tools showed that our segmentation algorithm tends to produce high-quality results that closely follow the visible boundaries of the neuronal somas (see Figure 4). It achieved superior results relative to the iterative thresholding method, which was prone to under-flooding or over-flooding depending on its input parameters and the initial shape of the soma. This is because if a segmentation uniformly over-floods out of its soma, its center of mass will remain consistent, meaning the over-flooding will avoid being detected. Similarly, if a soma has an odd shape, its center of mass movement will be captured too early, meaning the soma will not be allowed to be fully segmented. Global Otsu thresholding also yielded inaccurate results, prone to under-flooding somas whose intensities are lower than average, and over-flooding those somas whose intensities are higher than average, reflecting the fact that a global threshold is too generic to describe the specific intensities of each soma.

**Fig. 4.**
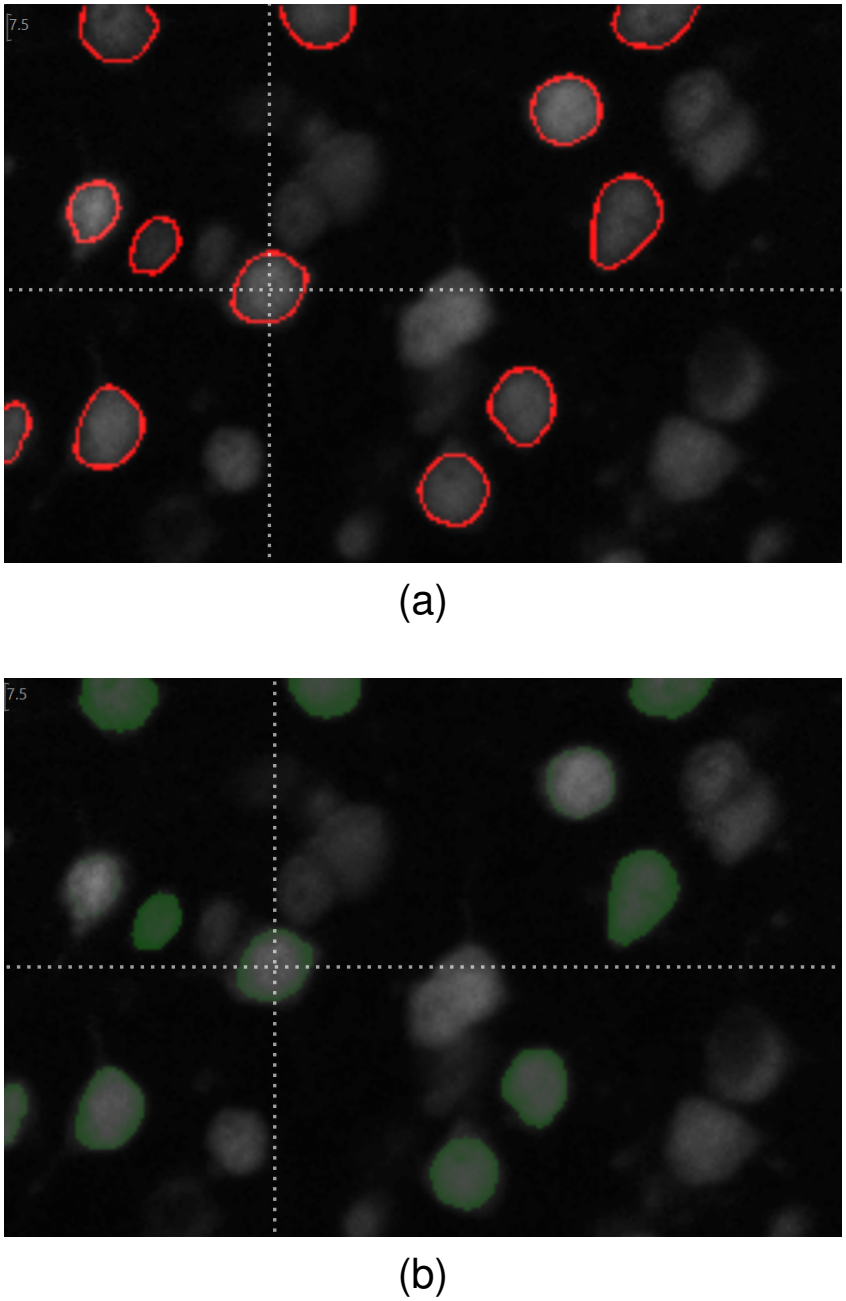
Segmented Somas in data obtained at a magnification of 9× . The somas are viewed with the 2D viewer in Vaa3D, and image contrast is increased to 67% during visualization to increase soma visibility, but is not changed during segmentation. (a) Results are displayed with an outline of the segmentation obtained using the Sobel filter. (b) The binary segmentation is transparently overlaid over the original image.

Overall, the visual tools prove critical in determining which specific somas produced poor results and guiding the user through manual threshold corrections. Most notably, the overlay with thin lines effectively assessed segmentation boundaries and their accuracy. These results suggest that a semi-automatic approach can significantly improve the speed and usability of labeling methods compared to purely manual ones while striking a balance between fidelity and quantity of generated data.

The results of the PCA of the somas in the binary segmentation can also be visualized to confirm that the components correctly capture each soma’s characteristic shape. Figure 5 shows these results in 3D, depicting their accuracy through visual inspection.

**Fig. 5.**
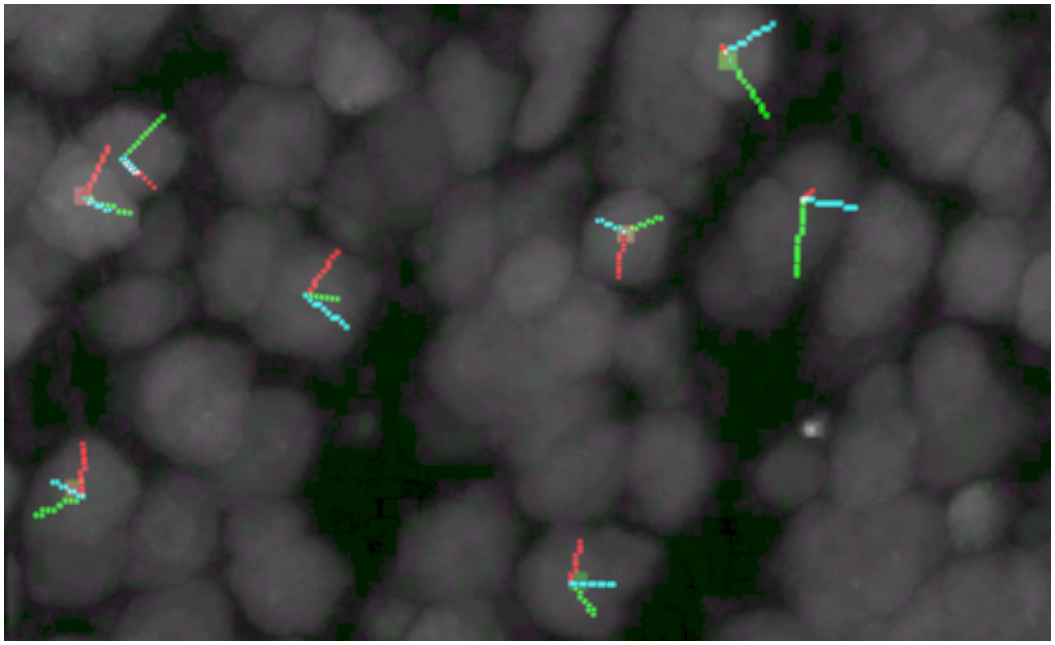
PCA Results performed on the segmentation of somas at a magnification of 9*×*. The somas are viewed with the 3D viewer in Vaa3D.

### C. Probabilistic Model of Soma Shape

The characteristic shapes of somas in different brain parts are captured within the probabilistic model of soma shape. This model is returned as one file that is part of the segmentation results and can be visualized as a heatmap using the plugin. As shown in Figure 6, qualitative inspection of results demonstrates that the model effectively depicts a soma’s characteristic shape.

**Fig. 6.**
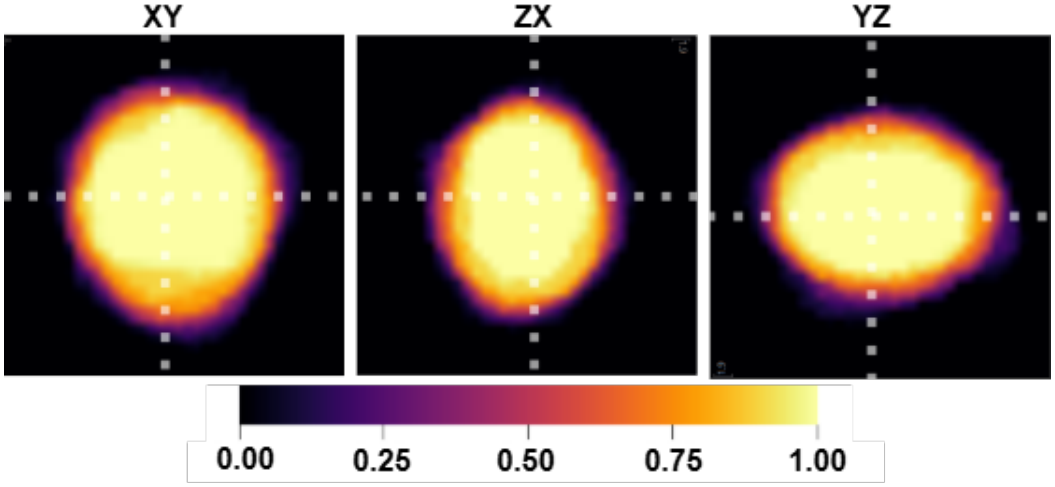
The probabilistic model demonstrates the characteristic shape of a neuronal soma. Axis labels are shown with the horizontal first and the vertical second. The Y axis has the longest principal component, the X axis has the second longest, and the Z axis has the shortest. Results were obtained by segmenting 10 neuronal somas of the 9× dataset in the same region of the brain used for Figure 4.

### D. Simulation

Quantitative evaluation of the effectiveness of the data augmentation process will be performed in future work, where a deep learning model will be trained with and without the simulated data obtained through this pipeline. The data augmentation’s performance evaluation will depend on the impact of the simulated data on the model’s accuracy.

Qualitative metrics used to evaluate simulation quality include how similar the simulation is to the original image, how realistic the synthetic image and its deformations are, and how accurate the segmentation is. As seen in Figure 7, the visual inspection tools integrated into Vaa3D show that our simulation produce high-quality synthetic images that are realistic variants of the original image. The segmentation label results also closely follow the performance obtained from our segmentation algorithm. However, the numerical errors that result from soma segmentation as previously discussed are visible in the slightly grainier outline of each soma in the synthetic image in Figure 7c. More advanced methods for addressing these numerical errors can be pursued in future work.

**Fig. 7.**
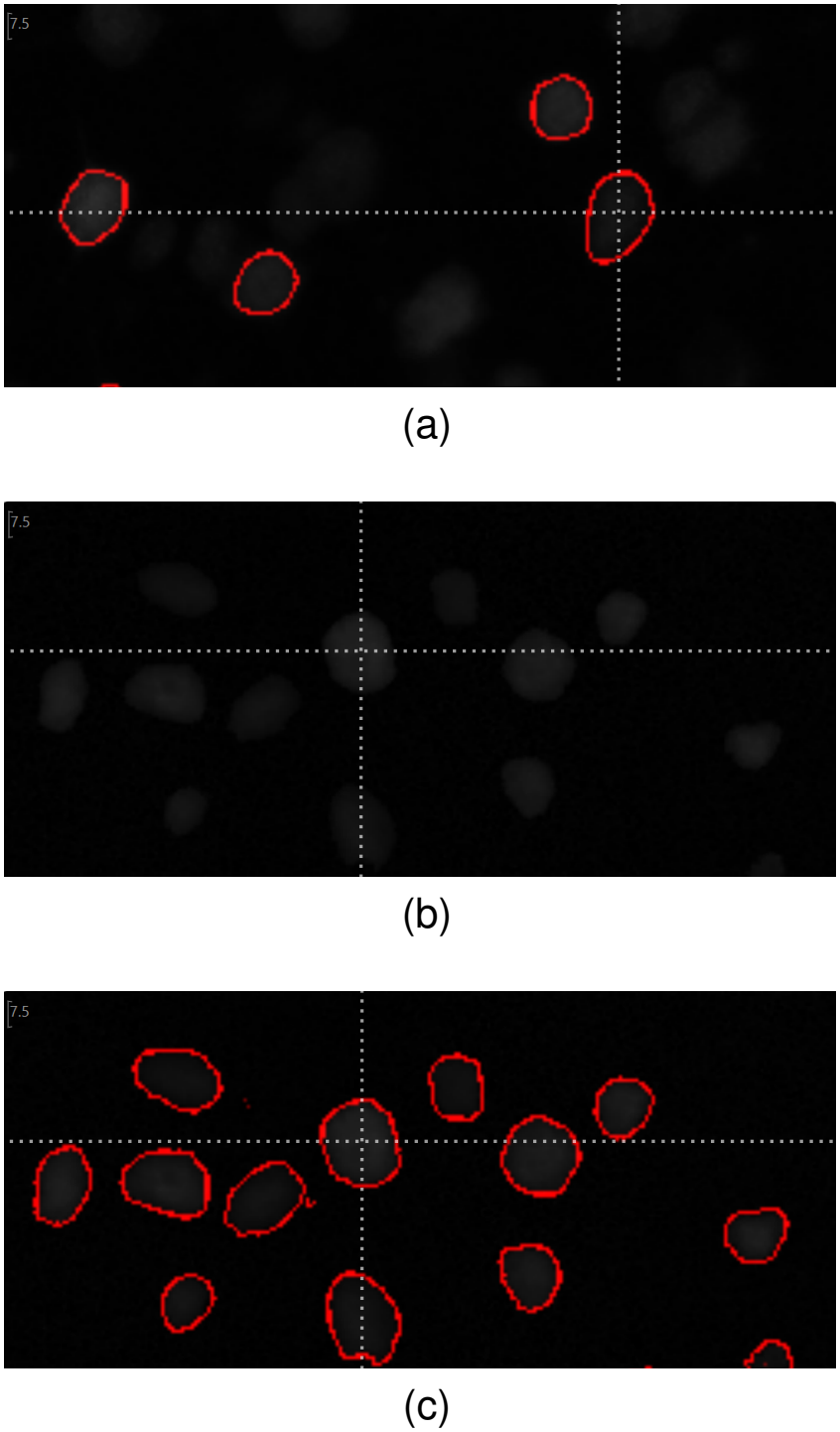
Comparison of real and synthetic soma data, with no change to image contrast to preserve the original image intensity. Synthetic data was generated based on 10 labeled neuronal somas imaged with magnification 9× . (a) The original image, including four segmented somas. (b) The simulated image. (c) The segmentation of the simulated image

Due to current limitations of the neuronal soma simulation pipeline, the resulting soma intensity, texture, shape, and segmentation accuracy heavily depend on the prior segmentation’s performance, as synthetic somas are reoriented and positioned copies of real somas, using a probabilistic model of neuron shapes. Future work to resolve these limitations can improve the simulation’s performance, as well as investigating more advanced techniques to generate synthetic shapes and intensities. This work is intended to serve as an initial basis for a simulation pipeline and is made open-source to enable improvements to its performance.

## V. Conclusion

The functions in the Machine Intelligence for Neuron Detection and Segmentation (MINDS) plugin for the open-source software platform Visually-assisted Analysis 3D (Vaa3d) successfully deliver a platform to semi-automatically segment neuronal somas, as well as to model their characteristic shape and to simulate ground truth data. The plugin integrates with Vaa3D and TeraFly, supporting the seamless navigation and analysis of terabytes of data, ensuring scalability towards large image datasets. The platform’s primary goal is to enable semi-automatic labeling and analyzing neuronal somas from a 3D brain volume obtained through LSFM. However, its results apply to cell segmentation of any intensity-based microscopy data.

The platform focuses on segmenting somas that can be isolated within a sphere around their center of mass. A local Ostu thresholding algorithm is used to determine the optimal threshold defining the boundary of each soma, and a 3D region-growing algorithm starting from a soma’s center of mass is applied to segment each one. Segmentation results can be analyzed qualitatively and quantitatively through the plugin’s tools, including an outline and overlay of segmentation results against the original image. These tools indicate a high-performing segmentation. A soma’s segmentation can be corrected by manually overriding the intensity threshold that defines its boundary. Overall, these features promise to significantly shorten the time researchers spend obtaining ground truth data by labeling every voxel in the image as part of a soma or the background, creating data that can be used for training deep learning pipelines for soma segmentation more accessible.

The soma segmentation plugin also produces PCA results for each segmented soma, providing insight into their orientation and characteristic shape. The plugin supports visualizing these principal components to confirm their accuracy and uses them to build a probabilistic model of soma shape. This provides further insight into somas’ characteristics and lays the foundation to create advanced simulations of neuronal somas in the brain.

Along this line, the plugin also provides a soma simulation pipeline, which creates synthetic ground truth data by repositioning and reorienting segmented somas in corresponding synthetic binary and realistic images, and uses the probabilistic model of soma shape and some simple probabilistic analysis to realistically deform the soma’s shape, and make its intensities synthetic as well. Qualitative inspection of simulated data reveals realistic images whose binary segmentation and intensities seem to match the accuracy of the initial segmented data, with some minor distortion due to numerical errors. The plugin supports multiple simulations of any image in the brain, promising to be an effective data augmentation technique to increase the quality and diversity of labeled data. We note that simulation creates data that can be used for training and ground truth for evaluating the performance of learning methods, because the user has all the information on each simulated soma. We expect that deep learning training can be pursued on these simulated data first, then transfer learning can be pursued for training with real data labeled by human raters. We expect that such a protocol will make use of our simulated data to improve performance.

The platform’s current implementation also has several limitations. It focuses on segmenting somas sufficiently isolated to be bound by a spherical marker. Isolating each soma is necessary for accurate analysis of soma shape and characteristics. Therefore, we expect that our tool will not perform as well in high neuron-density regions of the brain. Our future work could improve performance by developing extensions that allow the isolation of somas even in regions of high density.

Future work on soma simulation could explore advanced methods for deforming soma shapes based on the probabilistic model, as well as for creating synthetic soma intensity within this shape. This will improve the simulations’ production of realistic images, and create more accurate soma shapes for deep learning models to be trained on.

The ultimate goal remains to integrate deep learning models into this platform for automatic soma segmentation. The models can leverage both the semi-automatically labeled and the simulated ground truth data. This automation of the entire process of soma segmentation with increased accuracy would offer an accessible and cost-effective solution for all researchers.

## References

[1] D. S. Richardson, W. Guan, K. Matsumoto, C. Pan, K. Chung, A. Ertürk, H. R. Ueda, and J. W. Lichtman, “Tissue clearing,” Nature Reviews Methods Primers, vol. 1, no. 1, p. 84, 2021, publisher: Nature Publishing Group UK London.

[2] L. Ma, Q. Zhong, Y. Wang, X. Yang, and Q. Du, “A 3D semantic segmentation network for accurate neuronal soma segmentation,” Journal of Innovative Optical Health Sciences, p. 2450018, 2024, iSBN: 1793-5458 Publisher: World Scientific.

[3] X. Liu, B. Hu, M. Li, W. Huang, Y. Zhang, and Z. Xiong, “A soma segmentation benchmark in full adult fly brain,” in Proceedings of the IEEE/CVF Conference on Computer Vision and Pattern Recognition, 2023, pp. 7402–7411.

[4] R. Xiao, L. Zhu, J. Liao, X. Wu, H. Gong, J. Huang, P. Li, B. Sheng, and S. Chen, “DDeep3M+: adaptive enhancement powered weakly supervised learning for neuron segmentation,” Neurophotonics, vol. 10, no. 3, pp. 035 003–035 003, 2023, iSBN: 2329-423X Publisher: Society of Photo-Optical Instrumentation Engineers.

[5] X. Wei, Q. Liu, M. Liu, Y. Wang, and E. Meijering, “3D soma detection in large-scale whole brain images via a two-stage neural network,” IEEE Transactions on Medical Imaging, vol. 42, no. 1, pp. 148–157, 2022, iSBN: 0278-0062 Publisher: IEEE.

[6] M. Dong, D. Liu, Z. Xiong, C. Yang, X. Chen, Z.-J. Zha, G. Bi, and F. Wu, “3D CNN-based soma segmentation from brain images at single-neuron resolution,” in 2018 25th IEEE International Conference on Image Processing (ICIP). IEEE, 2018, pp. 126–130.

[7] O. Ronneberger, P. Fischer, and T. Brox, “U-Net: Convolutional Networks for Biomedical Image Segmentation,” 2015, eprint: 1505.04597. [Online]. Available: https://arxiv.org/abs/1505.04597

[8] P. A. Santi, “Light sheet fluorescence microscopy: a review,” Journal of Histochemistry & Cytochemistry, vol. 59, no. 2, pp. 129–138, 2011, publisher: SAGE Publications Sage CA: Los Angeles, CA.

[9] H. Peng, M. Hawrylycz, J. Roskams, S. Hill, N. Spruston, E. Meijering, and G. A. Ascoli, “BigNeuron: large-scale 3D neuron reconstruction from optical microscopy images,” Neuron, vol. 87, no. 2, pp. 252–256, 2015, publisher: Elsevier.

[10] L. Zhang, L. Huang, Z. Yuan, Y. Hang, Y. Zeng, K. Li, L. Wang, H. Zeng, X. Chen, H. Zhang, J. Xi, D. Chen, Z. Gao, L. Le, J. Chen, W. Ye, L. Liu, Y. Wang, and H. Peng, “Collaborative augmented reconstruction of 3D neuron morphology in mouse and human brains,” Nature Methods, vol. 21, no. 10, pp. 1936–1946, Oct. 2024. [Online]. Available: 10.1038/s41592-024-02401-8

[11] Y. Liu, S. Jiang, Y. Li, S. Zhao, Z. Yun, Z.-H. Zhao, L. Zhang, G. Wang, X. Chen, L. Manubens-Gil, Y. Hang, Q. Gong, Y. Li, P. Qian, L. Qu, M. Garcia-Forn, W. Wang, S. De Rubeis, Z. Wu, P. Osten, H. Gong, M. Hawrylycz, P. Mitra, H. Dong, Q. Luo, G. A. Ascoli, H. Zeng, L. Liu, and H. Peng, “Neuronal diversity and stereotypy at multiple scales through whole brain morphometry,” Nature Communications, vol. 15, no. 1, p. 10269, Nov. 2024. [Online]. Available: 10.1038/s41467-024-54745-6

[12] G. Lin, U. Adiga, K. Olson, J. F. Guzowski, C. A. Barnes, and B. Roysam, “A hybrid 3D watershed algorithm incorporating gradient cues and object models for automatic segmentation of nuclei in confocal image stacks,” Cytometry Part A, vol. 56A, no. 1, pp. 23–36, 2003, eprint: https://onlinelibrary.wiley.com/doi/pdf/10.1002/cyto.a.10079. x[Online]. Available: https://onlinelibrary.wiley.com/doi/abs/10.1002/cyto.a.10079

[13] Christ, “Segmentation of Medical Image using Clustering and Watershed Algorithms,” American Journal of Applied Sciences, vol. 8, no. 12, pp. 1349–1352, Dec. 2011. [Online]. Available: http://www.thescipub.com/abstract/10.3844/ajassp.2011.1349.1352

[14] X. Li, Z. Zhou, P. Keller, H. Zeng, T. Liu, and H. Peng, “Interactive exemplar-based segmentation toolkit for biomedical image analysis,” in 2015 IEEE 12th International Symposium on Biomedical Imaging (ISBI), 2015, pp. 168–171.

[15] L. Nichele, V. Persichetti, M. Lucidi, and G. Cincotti, “Quantitative evaluation of ImageJ thresholding algorithms for microbial cell counting,” OSA Continuum, vol. 3, no. 6, p. 1417, Jun. 2020. [Online]. Available: https://opg.optica.org/abstract.cfm?URI=osac-3-6-1417

[16] H. Peng, Z. Ruan, F. Long, J. H. Simpson, and E. W. Myers, “V3D enables real-time 3D visualization and quantitative analysis of large-scale biological image data sets,” Nature Biotechnology, vol. 28, no. 4, pp. 348–353, Apr. 2010. [Online]. Available: 10.1038/nbt.1612

[17] H. Peng, A. Bria, Z. Zhou, G. Iannello, and F. Long, “Extensible visualization and analysis for multidimensional images using Vaa3D,” Nature protocols, vol. 9, no. 1, pp. 193–208, 2014, iSBN: 1754-2189 Publisher: Nature Publishing Group UK London.

[18] H. Peng, J. Tang, H. Xiao, A. Bria, J. Zhou, V. Butler, Z. Zhou, P. T. Gonzalez-Bellido, S. W. Oh, J. Chen, A. Mitra, R. W. Tsien, H. Zeng, G. A. Ascoli, G. Iannello, M. Hawrylycz, E. Myers, and F. Long, “Virtual finger boosts three-dimensional imaging and microsurgery as well as terabyte volume image visualization and analysis,” Nature Communications, vol. 5, no. 1, p. 4342, Jul. 2014. [Online]. Available: 10.1038/ncomms5342

[19] A. Bria, G. Iannello, L. Onofri, and H. Peng, “TeraFly: real-time three-dimensional visualization and annotation of terabytes of multidimensional volumetric images,” Nature Methods, vol. 13, no. 3, pp. 192–194, Mar. 2016. [Online]. Available: 10.1038/nmeth.3767

[20] A. Bria, G. Iannello, and H. Peng, “An open-source VAA3D plugin for real-time 3D visualization of terabyte-sized volumetric images,” in 2015 IEEE 12th International Symposium on Biomedical Imaging (ISBI), 2015, pp. 520–523.

[21] J. Liang, L. Deng, S. Chen, Y. Wang, Z. Ruan, and L. Zhang, “Vaa3D-x for cross-platform teravoxel-scale immersive exploration of multidimensional image data,” Bioinformatics, vol. 39, no. 1, p. btac794, 2023, iSBN: 1367-4811 Publisher: Oxford University Press.

[22] L. Cai, J. Gao, and D. Zhao, “A review of the application of deep learning in medical image classification and segmentation,” Annals of Translational Medicine, vol. 8, no. 11, pp. 713–713, Jun. 2020. [Online]. Available: http://atm.amegroups.com/article/view/36944/html

[23] Ç içek, A. Abdulkadir, S. S. Lienkamp, T. Brox, and O. Ronneberger, “3D U-Net: Learning Dense Volumetric Segmentation from Sparse Annotation,” 2016, eprint: 1606.06650. [Online]. Available: https://arxiv.org/abs/1606.06650

[24] V. Badrinarayanan, A. Kendall, and R. Cipolla, “SegNet: A Deep Convolutional Encoder-Decoder Architecture for Image Segmentation,” IEEE Transactions on Pattern Analysis and Machine Intelligence, vol. 39, no. 12, pp. 2481–2495, 2017.

[25] K. Sun, B. Xiao, D. Liu, and J. Wang, “Deep High-Resolution Representation Learning for Human Pose Estimation,” 2019, eprint: 1902.09212. [Online]. Available: https://arxiv.org/abs/1902.09212

[26] O. Oktay, J. Schlemper, L. L. Folgoc, M. Lee, M. Heinrich, K. Misawa, K. Mori, S. McDonagh, N. Y. Hammerla, B. Kainz, B. Glocker, and D. Rueckert, “Attention U-Net: Learning Where to Look for the Pancreas,” 2018, eprint: 1804.03999. [Online]. Available: https://arxiv.org/abs/1804.03999

[27] X. Xiao, S. Lian, Z. Luo, and S. Li, “Weighted Res-UNet for High-Quality Retina Vessel Segmentation,” in 2018 9th International Conference on Information Technology in Medicine and Education (ITME), 2018, pp. 327–331.

[28] Z.-L. Ni, G.-B. Bian, X.-H. Zhou, Z.-G. Hou, X.-L. Xie, C. Wang, Y.-J. Zhou, R.-Q. Li, and Z. Li, “RAUNet: Residual Attention U-Net for Semantic Segmentation of Cataract Surgical Instruments,” 2019, eprint: 1909.10360. [Online]. Available: https://arxiv.org/abs/1909.10360

[29] P. Chlap, H. Min, N. Vandenberg, J. Dowling, L. Holloway, and A. Haworth, “A review of medical image data augmentation techniques for deep learning applications,” Journal of Medical Imaging and Radiation Oncology, vol. 65, no. 5, pp. 545–563, Aug. 2021. [Online]. Available: https://onlinelibrary.wiley.com/doi/10.1111/1754-9485.13261

[30] C. Shorten and T. M. Khoshgoftaar, “A survey on Image Data Augmentation for Deep Learning,” Journal of Big Data, vol. 6, no. 1, p. 60, Dec. 2019. [Online]. Available: https://journalofbigdata.springeropen.com/articles/10.1186/s40537-019-0197-0

[31] S. Yang, W. Xiao, M. Zhang, S. Guo, J. Zhao, and F. Shen, “Image Data Augmentation for Deep Learning: A Survey,” 2022. [Online]. Available: https://arxiv.org/abs/2204.08610

[32] J. Corral Acero, E. Zacur, H. Xu, R. Ariga, A. Bueno-Orovio, P. Lamata, and V. Grau, “SMOD - Data Augmentation Based on Statistical Models of Deformation to Enhance Segmentation in 2D Cine Cardiac MRI,” in Functional Imaging and Modeling of the Heart, Y. Coudiére, V. Ozenne, E. Vigmond, and N. Zemzemi, Eds. Cham: Springer International Publishing, 2019, vol. 11504, pp. 361–369. [Online]. Available: http://link.springer.com/10.1007/978-3-030-21949-939

[33] D. Karimi, G. Samei, C. Kesch, G. Nir, and S. E. Salcudean, “Prostate segmentation in MRI using a convolutional neural network architecture and training strategy based on statistical shape models,” International Journal of Computer Assisted Radiology and Surgery, vol. 13, no. 8, pp. 1211–1219, Aug. 2018. [Online]. Available: http://link.springer.com/10.1007/s11548-018-1785-8

[34] Z. Tang, K. Chen, M. Pan, M. Wang, and Z. Song, “An Augmentation Strategy for Medical Image Processing Based on Statistical Shape Model and 3D Thin Plate Spline for Deep Learning,” IEEE Access, vol. 7, pp. 133 111–133 121, 2019. [Online]. Available: https://ieeexplore.ieee.org/document/8835020/

[35] R. Bhalodia, A. Goparaju, T. Sodergren, R. Whitaker, A. Morris, E. Kholmovski, N. Marrouche, J. Cates, and S. Elhabian, “Deep Learning for End-to-End Atrial Fibrillation Recurrence Estimation,” Dec. 2018. [Online]. Available: http://www.cinc.org/archives/2018/pdf/CinC2018-160.pdf

[36] “vaa3d tools/released plugins/v3d plugins/cellseg quickfind at master · Vaa3D/vaa3d tools.” [Online]. Available: https://github.com/Vaa3D/vaa3dtools/tree/master/releasedplugins/v3dplugins/cellsegquickfind

[37] B. Charmouti, A. K. Junoh, W. Azman, W. Z. A. Wan Muhamad, M. Mansor, Z. Hasan, and M. Mashor, “Extended Median Filter For Salt and Pepper Noise In Image,” International Journal of Applied Engineering Research, vol. 12, pp. 12 914–12 918, Jan. 2017.

[38] O. Vincent and O. Folorunso, “A descriptive algorithm for sobel image edge detection,” 01 2009.

[39] G. Bradski, “The opencv library.” Dr. Dobb’s Journal: Software Tools for the Professional Programmer, vol. 25, no. 11, pp. 120–123, 2000.

[40] J. E. Bresenham, “Algorithm for computer control of a digital plotter,” in Seminal graphics: pioneering efforts that shaped the field, 1998, pp. 1–6.

[41] O. Rainio, J. Hällilä, J. Teuho, and R. Klén, “Methods for estimating full width at half maximum,” Signal, Image and Video Processing, vol. 19, no. 4, p. 289, Feb. 2025. [Online]. Available: 10.1007/s11760-025-03820-6

[42] N. Otsu, “A Threshold Selection Method from Gray-Level Histograms,” IEEE Transactions on Systems, Man, and Cybernetics, vol. 9, no. 1, pp. 62–66, 1979.

[43] M. Abdolhoseini, M. G. Kluge, F. R. Walker, and S. J. Johnson, “Neuron Image Synthesizer Via Gaussian Mixture Model and Perlin Noise,” in 2019 IEEE 16th International Symposium on Biomedical Imaging (ISBI 2019), 2019, pp. 530–533.

[44] X. Ruan, M. Mueller, G. Liu, F. Görlitz, T.-M. Fu, D. E. Milkie, J. L. Lillvis, A. Kuhn, J. Gan Chong, J. L. Hong, C. Y. A. Herr, W. Hercule, M. Nienhaus, A. N. Killilea, E. Betzig, and S. Upadhyayula, “Image processing tools for petabyte-scale light sheet microscopy data,” Nature Methods, vol. 21, no. 12, pp. 2342–2352, Dec. 2024. [Online]. Available: 10.1038/s41592-024-02475-4

[45] G. E. P. Box and M. E. Muller, “A Note on the Generation of Random Normal Deviates,” Annals of Mathematical Statistics, vol. 29, pp. 610–611, 1958. [Online]. Available: https://api.semanticscholar.org/CorpusID:119971394

[46] C. Karpfinger, “The QR-Decomposition of a Matrix,” in Calculus and Linear Algebra in Recipes: Terms, phrases and numerous examples in short learning units, C. Karpfinger, Ed. Berlin, Heidelberg: Springer Berlin Heidelberg, 2022, pp. 189–199. [Online]. Available: 10.1007/978-3-662-65458-319

[47] G. Guennebaud, B. Jacob, and others, “Eigen v3,” 2010. [Online]. Available: http://eigen.tuxfamily.org

